# Cytomegalovirus-induced T cell responses accelerate Alzheimer’s disease progression in mice

**DOI:** 10.1101/2025.03.28.645902

**Authors:** Morgan Marsden, James E McLaren, Ryan J Bevan, Daisy Penn-Ripley, Michelle Somerville, Sarah N Lauder, Awen Gallimore, David A Price, Kelly L Miners, Kristin Ladell, Florian A Siebzehnrubl, Timothy R Hughes, Ian R Humphreys, Mathew Clement

## Abstract

Infections have long been implicated as causative factors in Alzheimer’s disease (AD). Multiple studies have further suggested a key role for herpesviruses, such as cytomegalovirus (CMV). Using transgenic 3xTg-AD mice, we demonstrate that systemic infection with the β-herpesvirus murine cytomegalovirus (MCMV) accelerates the development of cognitive decline, tauopathy, and synaptic loss in the hippocampus, all of which are key features of AD. Accelerated disease progression after infection was associated with substantial lymphocyte infiltration into the brain dominated by CD8^+^ T cells specific for MCMV. Moreover, T cell receptor analyses revealed that these responses were clonally diverse, suggesting that multiple viral antigens were targeted in the brain during chronic infection with MCMV. T cell depletion during virus chronicity rescued infection-induced cognitive decline. In addition, antiviral drug treatment reduced lymphocytic infiltrates in the brain and reversed cognitive decline, suggesting potential clinical utility. These data provide a mechanistic link between chronic viral infections and the development of AD.

## Introduction

Alzheimer’s disease (AD) is the most common form of dementia, with ∼50 million cases worldwide and a predicted ∼152 million cases by 2050 [1]. Although a number of genetic risk factors for both early- and late-onset AD have been identified (reviewed in: [2]), the environmental factors that increase the risk of AD development and/or progression are not fully understood.

The hypothesis that infectious diseases contribute to AD development was presented over a century ago by Alzheimer, Fischer and colleagues due to similarities in AD pathologies with syphilitic dementia [3, 4]. Since then, bacterial infections including *Borrelia* species have been implicated in AD (reviewed in: [5]). Importantly, numerous viruses ranging from influenza [6] to SARS-CoV-2 [7] have also been implicated in AD. Notably, chronic human herpesviruses (HHV) have received particular attention as risk factors in AD due to their neurotropism, and the presence of herpes simplex virus (HSV) [8, 9], HHV6-A and HHV-7 [10] DNA in the brains of AD sufferers has been reported. Furthermore, the recombinant shingles vaccine targeting the herpesvirus varicella-zoster virus (VZV) reduces risk of dementia development in the 6 years post-vaccination [11]. Although not classically considered as neurotropic in immune competent individuals, the γ-herpesvirus human cytomegalovirus (HCMV) has also been associated with increased risk of AD development and accelerated cognitive decline [12–15]. Thus, substantial evidence implies that herpesviruses play a role in promoting AD progression. Despite these associations, however, the cellular and molecular mechanisms linking viral infections to AD development are poorly understood. Identifying the biological processes that are induced or exacerbated by viruses during AD and defining how they contribute to disease development may lead to new treatment strategies for patients.

How the immune system, including T cells, contributes to AD development has been a focus of interest [16]. Studies of AD patients have reported the presence of T cells in the hippocampus, leptomeninges and cerebrospinal fluid [17–19], with similar findings reported in mouse models [19, 20]. AD neuropathology is characterised by neuritic plaques and neurofibrillary tangles (NFTs) that develop due to amyloid-beta accumulation (Aγ) and tau phosphorylation, leading to synaptic loss and neuronal cell dysfunction. T cells associate with tau pathology in humans [17] and, in mice, drive neurodegeneration in tauopathy [21], suggesting a mechanistic link between T cells with brain damage and associated cognitive decline.

T cells are present in brains of human AD patients post-mortem and in mouse models of AD [16] whilst severe disease is associated with an elevated CD8^+^ T cell response in the hippocampus [17, 19, 22]. This raises the possibility that immune insult resulting in inflammatory T cell responses may contribute to disease progression. Given that induction of robust T cell responses is a key feature of the host immune response to bacterial and viral infections, it is possible that virus-induced T cells may exacerbate AD development. In accordance with this hypothesis, studies of T cell infiltrates in the cerebrospinal fluid (CSF) of AD patients revealed the presence of clonally expanded CD8^+^ T cells reactive to the gamma herpesvirus Epstein-Barr virus (EBV) [18]. There have been multiple studies that have implicated HCMV as a key contributor to AD onset [12, 14, 23, 24]. However, the role that HCMV and the key immune mechanisms associated with infection that contribute to AD is poorly understood. We used the well-characterised murine cytomegalovirus (MCMV) experimental model of herpesvirus infection, which has been demonstrated to limits blood brain barrier integrity and cognition after repeated infectious challenges *in vivo* [25], to investigate whether virus-induced immune responses participate in viral-associated disease progression. Using the 3xTg-AD mouse model, we now reveal a direct role for herpesvirus-induced T cells in driving infection-induced brain pathology and cognitive decline in AD.

## Results

### Chronic MCMV infection accelerates cognitive decline in 3xTg-AD mice

To investigate how herpesvirus infection impacts AD development, we first established an *in vivo* experimental model of viral infection of 3xTg-AD mice using MCMV as a well-characterised model of acute and chronic herpesvirus infection. 3xTg-AD mice exhibited comparable virus-induced weight loss to wild type C57BL/6 control mice during acute infection (Fig. S1A) and virus shedding at the peak time-point of viral persistence (∼day (d)33) in the salivary glands (Fig. S1B), suggesting control of viral infection was comparable in C57BL/6 and 3xTg-AD mice. Although MCMV can persist in brains of neonatal mice, systemic infection of immunocompetent adult mice does not lead to measurable infection of brain tissue [26, 27]. In accordance, we failed to detect viral DNA in whole brain tissue from chronically-infected C57BL/6 or 3xTg-AD mice (Fig. S1C).

To investigate whether MCMV infection influences AD development, 3-4 month old female 3xTg-AD mice were infected with MCMV and cognitive ability was assessed. After 6 months of infection (∼10 months of age), mock-infected 3xTg-AD mice exhibited comparable cognition to C57BL/6 mice in both Novel Object Recognition (NOR) and T-maze alternation rate testing (Fig. 1 A-D). Importantly, virally-infected 3xTg-AD mice displayed impaired performance in NOR testing revealing reduced discrimination ratios (DR) by comparison with uninfected controls (Fig. 1A & B). Similarly, virus-infected 3xTg-AD mice displayed altered spatial working memory when compared to mock-infected 3xTg-AD and C57BL/6 mice (Fig. 1C), demonstrating that MCMV infection accelerates cognitive decline in 3xTg-AD mice.

**Figure 1.**
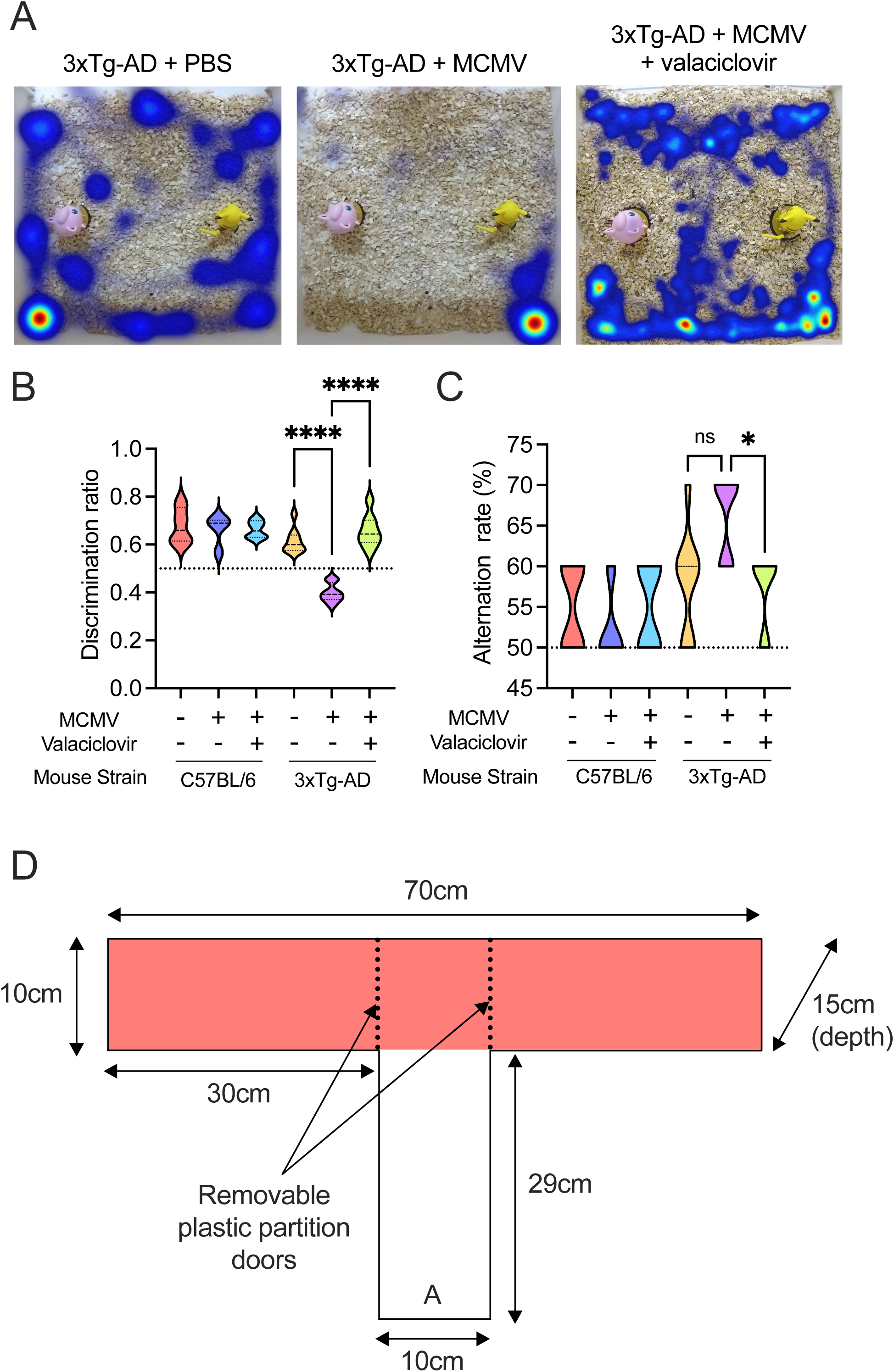
Chronic viral infection exacerbates cognitive decline in AD mice. (A-C) C57BL/6 or 3xTg-AD mice were infected with 5 × 10^4^ pfu MCMV or PBS i.p. Some groups of mice were also treated with 1mg/ml Valaciclovir Hydrochloride dissolved in animal drinking water continuously from day 60 post infection until experiment termination. At 9 months of age animal behaviour and cognitive assessment were measured using (A & B) novel object recognition (NOR) and (C) spontaneous alternation rate (AR). (A) Heatmap analysis tracking animal path and animal behaviour in 9-month-old 3xTg-AD mice after PBS treatment, or viral-infection +/-oral valaciclovir treatment. (B) Discrimination ratio (DR) determined post NOR assessment whereby impairment of cognition is shown by a DR below 0.5. (C) AR determined post T-maze test replicated 10 times/mouse/condition. AR of >60% per chosen open arm of test indicates loss or weakening of cognition. Data shown are representative of 2-5 separate experiments with 4 C57BL/6 and 10 3xTg-AD mice/group. Statistical significance was assessed using two-way ANOVA analysis with Tukey’s multiple comparisons test. p values are reported as follows: n.s., >0.05; *, ≤0.05; **, ≤0.01; ***, ≤0.001; and ****, ≤0.0001. (D) Schematic representation of T maze apparatus used for spontaneous alternation rate testing.

To ascertain whether chronic infection was a prerequisite for MCMV-induced cognitive decline, mice were orally treated (or not) with the antiviral drug valaciclovir hydrochloride, which exerts robust antiviral activity against MCMV [28], after the resolution of acute infection and mucosal persistence (d60 post-infection). Valaciclovir treatment during the chronic phase of infection completely reversed virus-driven cognitive decline when assessed by NOR and alternation rates (Fig. 1A-C). Thus, chronic MCMV infection accelerated cognitive decline in 3xTg-AD mice.

### MCMV infection induces neuronal damage and tauopathy in 3xTg-AD mice

To understand the neuropathological mechanisms underpinning MCMV-induced cognitive decline in 3xTg-AD mice, we first investigated whether MCMV impacted synaptic plasticity through measurements of CA1 hippocampal dendritic spines. 9-to 10-month-old C57BL/6 or 3xTg-AD mice infected with MCMV or mock infected with PBS for 7 months were assessed. At this time, mock-infected 3xTg-AD mice exhibited a non-significant trend in total spine loss as compared to C57BL/6 controls (Fig. 2A & B). In contrast, MCMV-infection caused significant (p <0.0005) total spine loss in MCMV-infected 3xTg-AD mice as compared to infected C57BL/6 controls (Fig. 2A & B). Moreover, when comparing 3xTg-AD mice, MCMV infection led to a significant loss in spine density (p <0.005) as compared to mock-infected mice (Fig. 2A & B). Differential spine counts indicated that MCMV chronicity led to a preferential loss in stubby and mushroom spines (Fig. 2A & B). These dendritic spines are classified by sub-types including thin (learning and plasticity), mushroom (long-term memory storage and synaptic connections), and stubby (common in early development and may represent a transient less mature form) [29, 30]. Thus, these data support the hypothesis that MCMV-induced cognitive decline associated with hippocampal neuronal damage.

**Figure 2.**
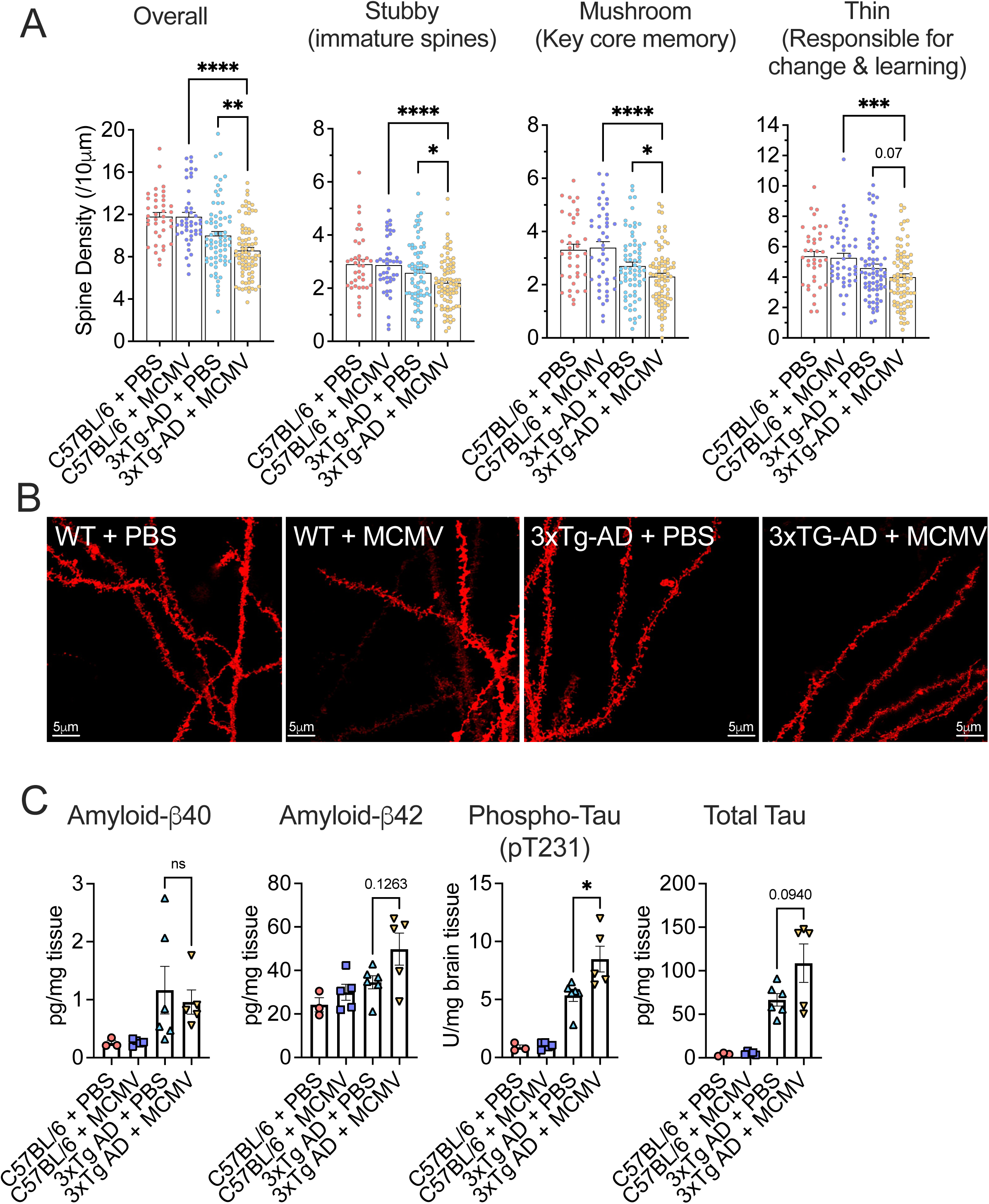
Chronic viral infection promotes loss of neuronal function. (A & B) DiOlistic labelling of CA1 hippocampal dendritic spines from C57BL/6 or 3xTg-AD mice infected with 5 × 10^4^ pfu MCMV or PBS i.p. At 9 months of age brains were harvested and the hippocampus was removed for DiOlistic labelling. (A) Overall and individual spine density morphology from each group is counted using Imaris software. (B) Representative confocal images of neuronal spines taken from the CA1 region of the hippocampus. (A) Data are shown as mean ± SEM; (A & B) from 4 C57BL/6 and 6 3xTg-AD mice per group with multiple data points from each mouse shown. Statistical significance was assessed using Welch’s t test. p values are reported as follows: n.s., >0.05; *, ≤0.05; **, ≤0.01; ***, ≤0.001; and ****, ≤0.0001. (C) Aγ40, Aγ42, phospho Tau (pT231) and total Tau concentration in brain homogenates of 9–10-month-old C57BL/6 or 3xTg-AD mice infected/not with 5 × 10^4^ pfu MCMV (i.p.) for 6-7 months measured by ELISA. Data shown are representative of 3-5 C57BL/6 and 5-6 3xTg-AD mice per group. Statistical significance was assessed using one-way ANOVA analysis with Tukey’s multiple comparisons test. p values are reported as follows: n.s., >0.05; *, ≤0.05; **, ≤0.01; ***, ≤0.001; and ****, ≤0.0001.

AD neuropathology is characterised by neuritic plaques and neurofibrillary tangles (NFTs) that develop due to amyloid-beta accumulation (Aγ) and tau phosphorylation, respectively. In 3xTg-AD mice (not infected), Aγ plaques have recently been shown to be only detectable from 12 months of age onwards in females [31]. However, upon MCMV infection of 3xTg-AD mice, we observed an elevation of Aγ42 (human) in brain homogenates at 9 months of age (Fig. 2C). Given that HCMV seropositivity associates with neurofibrillary tangles in individuals with AD [23] and MCMV induces Tau accumulation upon *in vitro* infection [32], we also measured the impact of MCMV infection on total and phosphorylated Tau (Tau-p) in brains. In brain homogenates from MCMV-infected 3xTg-AD mice this was associated with a significant increase in human Tau-p and an elevation of total human Tau 6 months post-infection (Fig. 2C). Thus, the accelerated neuronal damage observed in MCMV-infected 3xTg-AD mice associated with elevated Tau accumulation and, to a lesser extent, Aγ42 elevation.

### Robust brain-infiltrating CD8 T cell responses are induced by MCMV infection in 3xTg-AD mice

Given that we failed to detect measurable viral DNA in the brains of MCMV-infected mice, we reasoned that MCMV-induced neuronal damage and cognitive decline may be a consequence of virus-induced perturbation of host immune responses. MCMV infection in C57BL/6 mice induces robust T cell responses that, notably in the case of CD8^+^ T cells, increase in frequencies over time [33–35]. Given the association between T cells and tau pathology [17, 21], we investigated whether MCMV induced T cell infiltrations in brain tissue. We observed a significant accumulation of CD4^+^ and, more strikingly, CD8^+^ T cells in brains of infected 3xTg-AD mice (Fig. 3A). Virus-induced infiltrations of CD8^+^ T cells consisted of high frequencies of virus-specific T cells reactive to peptide epitopes derived from the viral IE3 and M38 proteins (Fig. 3B-D).

**Figure 3.**
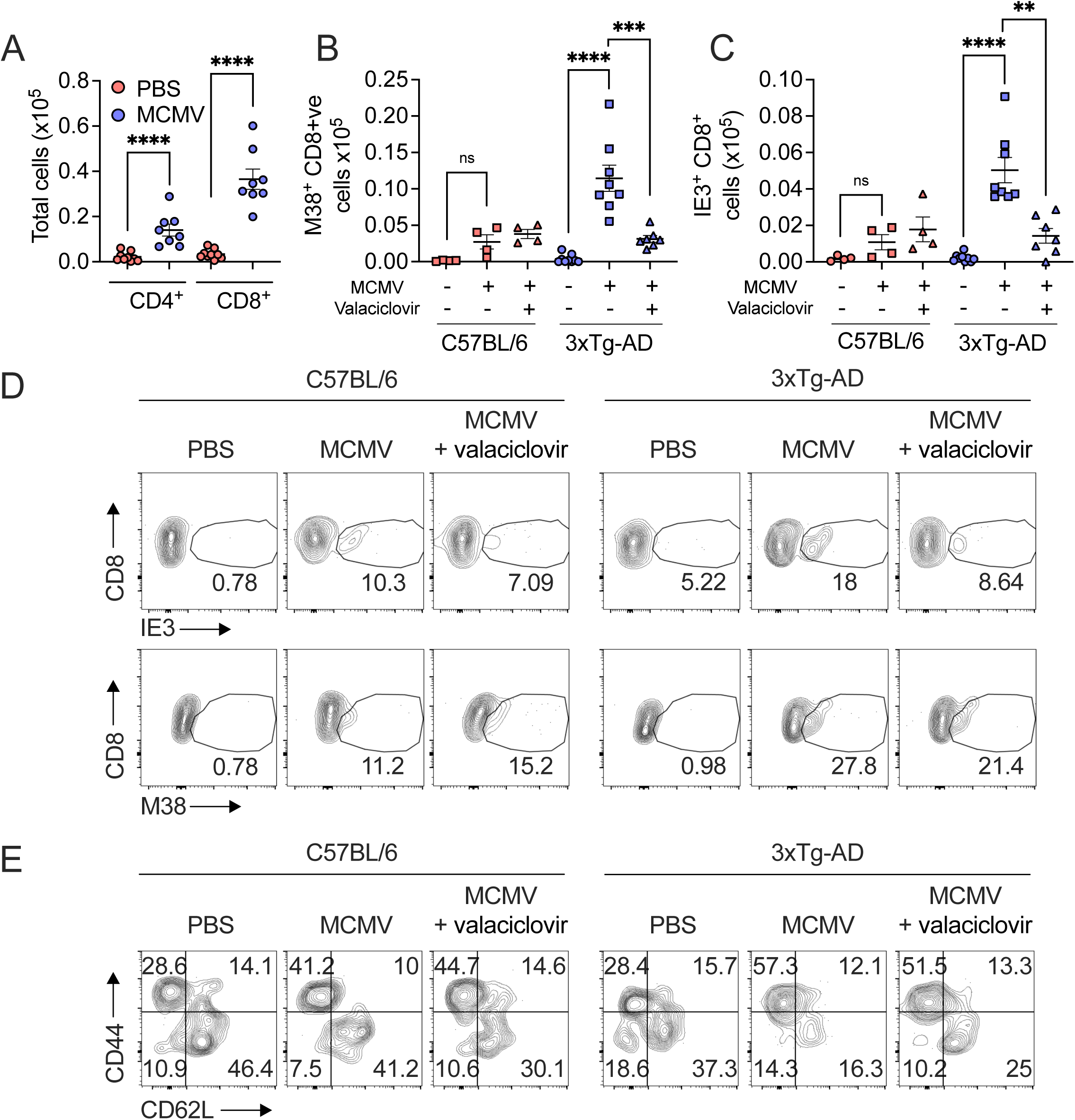
Virus infection induces increased T cell infiltrates in the brains of AD susceptible mice. C57BL/6 or 3xTg-AD mice were infected with 5 × 10^4^ MCMV or PBS i.p. Some groups of mice were also treated with 1 mg/ml Valaciclovir Hydrochloride dissolved in animal drinking water continuously from day 60 post infection until experiment termination. Leukocytes were harvested from the brains at 9 months of age (6 months post-infection). Total CD4^+^ and CD8^+^ T cells (A) and MCMV tetramer^+^ (M38 and IE3) CD8^+^ T cells (B-D) were quantified via flow cytometry. Statistical significance (A) was assessed using Mann– Whitney U test. p values are reported as follows: n.s., >0.05; *, ≤0.05; **, ≤0.01; ***, ≤0.001; and ****, ≤0.0001. Statistical significance (B & C) was assessed using two-way ANOVA analysis with Tukey’s multiple comparisons test. p values are reported as follows: n.s., >0.05; *, ≤0.05; **, ≤0.01; ***, ≤0.001; and ****, ≤0.0001. (E) Bivariant FACS plots showing percentage expression of CD44 versus CD62L in CD8^+^ T cells quantified in brains via flow cytometry. (A-E) Data are shown as mean ± SEM; n = 4 C57BL/6 and 9-10 3xTg-AD mice per group representing two independent experiments).

Phenotyping analysis revealed that MCMV infection induced the enrichment of CD8^+^ effector memory T cells (Tem) (CD44^+^/CD62L^-^) upon MCMV infection of C57BL/6 and, to a greater extent, 3xTg-AD mice (Fig. 3E). Interestingly, MCMV did not elicit a notable enrichment of tissue resident CD4^+^ or CD8^+^ T cells in either mouse strain (Fig. S2A). Thus, MCMV infection induced an enrichment of brain-infiltrating T cells that were predominantly effector memory T cells.

### Virus-specific CD8^+^ T cells with increased clonal diversity seed the brain during MCMV-accelerated cognitive decline

Site-specific immune responses can give rise to clonal CD8^+^ T cell expansions and specific bias within encoded T-cell receptors (TCRs) upon infection [36]. To determine whether MCMV induces T cell clonal expansions in brain tissue, we performed TCR deep sequencing of virus-specific (M38-reactive) CD8^+^ T cells in the brains and blood of mice. An assessment of murine TCR γ-chain variable (TRBV) region usage revealed a marked increase in TCR diversity in M38-specific TCRs in the brains of MCMV-infected 3xTg-AD mice, compared to C57BL/6 controls (Fig. 4A & B, Fig. S3A), consistent with the induction of a broad virus-specific CD8^+^ T cell response in the brains during AD development.

**Figure 4.**
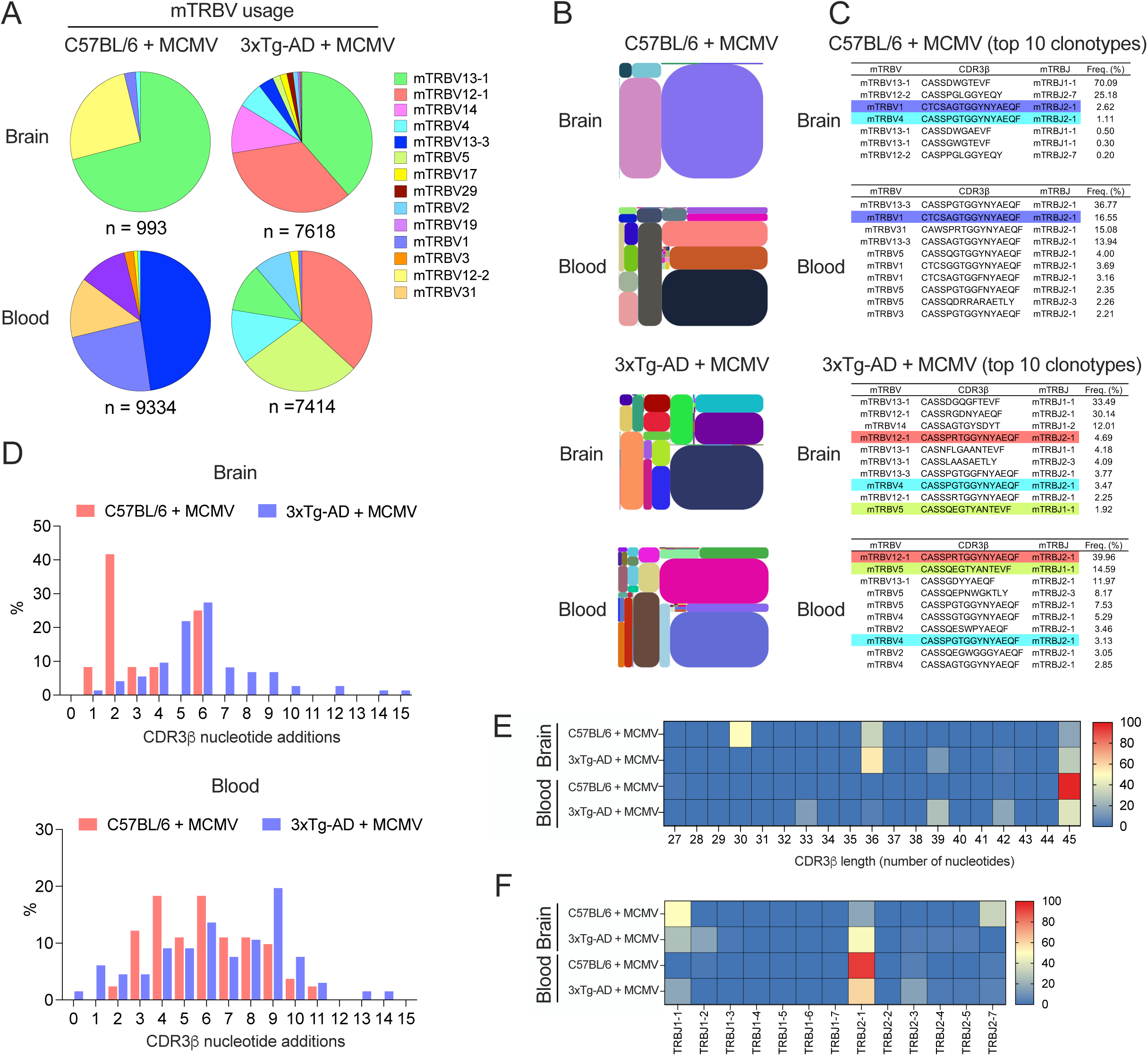
Viral-specific CD8^+^ T cells exhibit increased clonally diversity in AD susceptible mice upon cognitive decline. C57BL/6 or 3xTg-AD mice were infected with 5 × 10^4^ pfu MCMV (i.p.). Leukocytes were harvested from the brains and circulating blood at 6 months post-infection (9 months of age). (A) Concatenated frequency of murine TRBV (mTRBV) gene transcripts (key) and (B) tree maps of CDR3γ clonotype usage in relation to overall repertoire size of MCMV tetramer^+^ (M38) CD8^+^ T cells from the brain and blood of C57BL/6 or 3xTg-AD mice. Different colours represent different TCRs with size relating to total proportion of the defined repertoire. (C) TCR γ-chain amino acid sequences of M38-tetramer^+^ CD8^+^ T cells from the brain and blood of C57BL/6 or 3xTg-AD mice with top 10 most frequent clonotypes shown with coloured sequences representing clonotypes matching across tissue types and groups. (D) TCR convergence measured as the number of nucleotide sequences encoding amino acid-identical complementarity-determining region CDR3γ loops (x-axis) across the most prevalent clonotypes in the brain (upper panel) and the blood (lower panel) within M38-tetramer^+^ TCR γ-chains from the brain and blood of C57BL/6 or 3xTg-AD mice. (E & F) Heatmaps depicting distribution of (E) CDR3γ length and (F) TRBJ usage within M38-tetramer^+^ TCR γ-chains from the brain and blood of C57BL/6 or 3xTg-AD mice. All data shown are concatenated from M38-tetramer^+^ CD8^+^ T cells sorted from pooled C57BL/6 or 3xTg-AD mice (n = 5 per group, per tissue and is representative of two experiments)

Splenomegaly and peripheral blood perturbations are reported in 3xTg-AD mice [37], and we observed a non-significant increase in peripheral immune responses in 3xTg-AD versus C57BL/6 mice (Fig. S2B-D). However, TCR diversity within M38-specific CD8^+^ T cells in the blood was more comparable between 3xTg-AD and C57BL/6 mice (Fig. 4A & B, Fig. S3A & B), consistent with the conclusion that broad expansions of virus-specific CD8^+^ T cells in 3xTg-AD was restricted to brains of 3xTg-AD mice. Furthermore, whilst matching TCR clonotypes were identified in paired repertoires from the blood and brain of both C57BL/6 and 3xTg-AD mice, we detected increased TCR clonotype overlap in 3xTg-AD mice (Fig. 4A-C). These data are consistent with the conclusion that virus-specific T cells that are detectable in peripheral blood in 3xTg-AD mice are recruited to the brain during AD development.

The increase abundance of more prominent TCR clonal expansions in the brains of 3xTg-AD mice suggested that, whilst there was no detectable viral replication in the brains of viral-infected 3xTg-AD mice, these cells were more antigen-focussed than comparable cells in C57BL/6 mice. This was confirmed by the overall increase of nucleotide variants in viral-infected 3xTg-AD CD8^+^ T cells as compared the C57BL/6 mice (Fig. 4D). These variants are responsible for encoding the complementarity-determining region (CDR)3γ amino acid sequence and make direct contact with the pMHCI molecule giving rise to antigen-specific convergence [38]. This increase in nucleotide variants suggested higher TCR diversity through recombination in viral-infected 3xTg-AD mice (Fig. 4D & E) and is confirmed by an elevated Shannon entropy score in the 3xTg-AD as compared to the C57BL/6 mice (Fig. S3B). CDR3γ length and TCRBJ gene usage also differed between the blood and brain of viral-infected 3xTg-AD and C57BL/6 mice highlighting the unique increase in TCR diversity and clonotypic differences between the viral-specific T-cells in the brains of 3xTg-AD mice. Overall these data show that elevated recruitment to the brain of virus-specific CD8^+^ T cells with increased TCR clonal diversity occurs during MCMV infection in transgenic AD mice.

### Virus-induced T cells drive AD development in 3xTg mice

Histological analysis revealed that chronic MCMV infection of 3xTg-AD mice induced enrichment of CD8^+^ T cells in the hippocampus (Fig. 5A & G), where virus-associated neuronal damage occurs (Fig. 2A & B). Furthermore, despite the absence of detectable viral antigen at late time-points measured in these studies (Fig. S1C), we observed that concentrations of the T cell-activating cytokine IL-18, which can activate T cells in the absence of antigen [39], were higher in brain homogenates of MCMV-infected 3xTg-AD mice as compared to C57BL/6 controls (Fig. S4A). Consistent with the hypothesis that MCMV-induced T cells directly contribute to AD development, improved cognitive function in MCMV-infected 3xTg-AD mice that were treated with the antiviral drug valaciclovir (Fig. 1) was associated with a reduction in brain-infiltrating virus-specific T cell responses (Fig. 3B-D).

**Figure 5.**
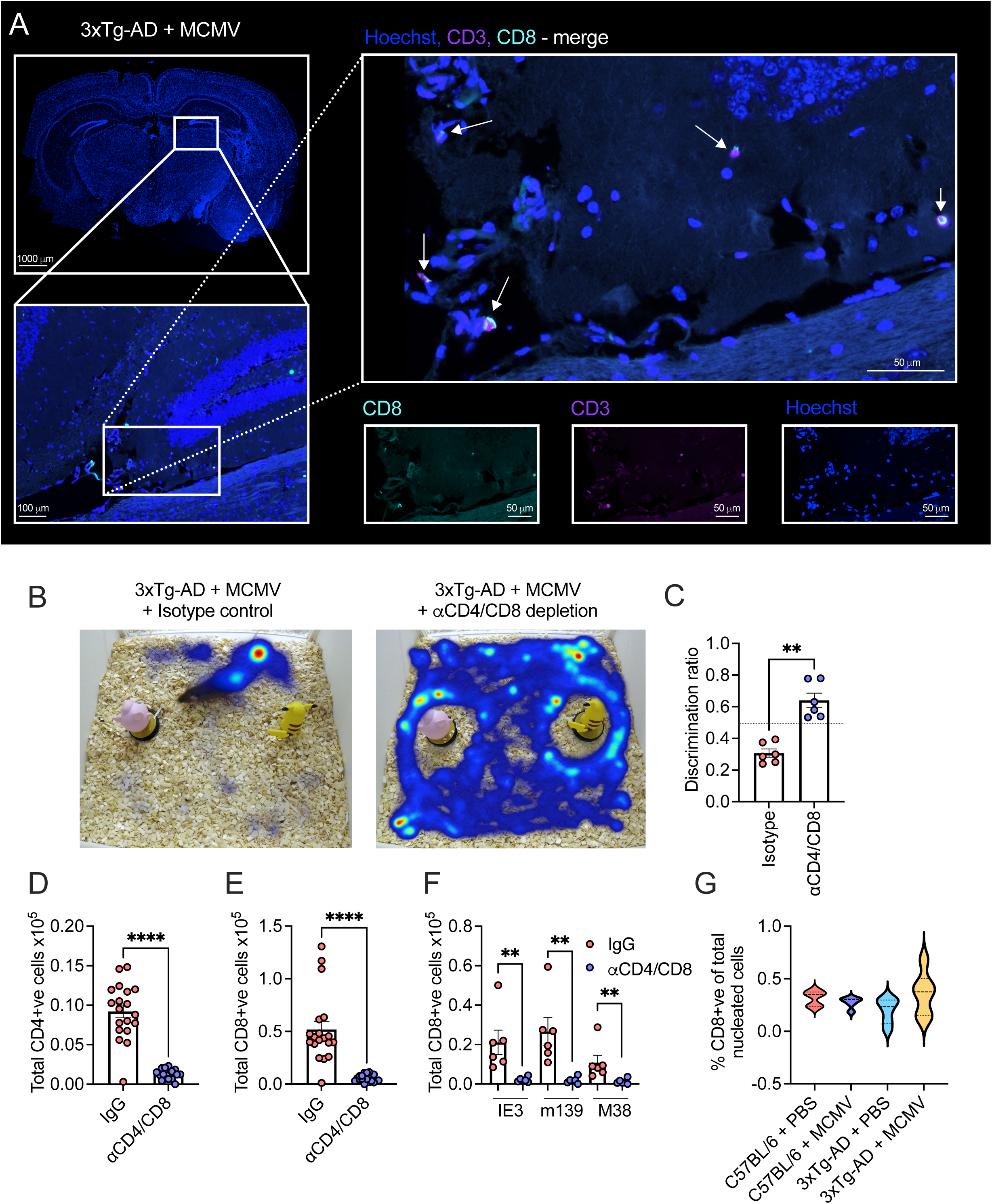
T cells drive CMV-induced cognitive decline. (A-G) C57BL/6 or 3xTg-AD mice were infected with 5 × 10^4^ pfu MCMV or PBS (i.p). Some mice were also treated with depleting anti-CD4 and anti-CD8 antibodies or isotype control. (A) Microscopy analysis of brain coronal section (upper left panel) of CD3^+^ (purple) /CD8^+^ (cyan) dual expressing T cells in the hippocampus (lower left and all right panels) of 9-month-old 3xTg-AD mice infected with 1 × 10^4^ pfu MCMV for 6 months. Data are shown as merged expression (upper right panel) with white arrows pointing to CD3^+^/CD8^+^ dual expressing T cells, and individual expression (bottom right panels). (B-F) At 9 months of age (6 months post infection) animal behaviour and cognitive assessment was measured using (B) novel object recognition (NOR) heatmaps with (C) Discrimination ratio (DR) determined post NOR assessment indicating impairment of cognition by a DR below 0.5. (D-F) Leukocytes were harvested from the brains at 9 months of age and accumulation of CD4^+^ T cells (D), CD8^+^ T cells (E) and MCMV tetramer^+^ (IE3^+^, m139^+^ and M38^+^) CD8^+^ T cells (F) were quantified via flow cytometry. (G) Percentage total CD3^+^/CD8^+^ cells (purple and cyan) from (A) of total nucleated cells (blue – Hoechst 33342) in the hippocampus of C57BL/6 or 3xTg-AD mice infected with 5 × 10^4^ pfu MCMV or PBS (i.p). Data are shown as mean ± SEM; (A-G) n = 5-6 C57BL/6 and 5-6 3xTg-AD mice per group representing two independent experiments). Statistical significance was assessed using Mann–Whitney U test. p values are reported as follows: n.s., >0.05; *, ≤0.05; **, ≤0.01; ***, ≤0.001; and ****, ≤0.0001.

Thus, to definitively test whether T cells contributed to MCMV-induced cognitive decline, we treated 3xTg-AD mice with anti-CD4 and anti-CD8 depleting antibody during MCMV chronicity. Treatment depleted CD4^+^ and CD8^+^ T cell responses in the brain at 9 months of age (6 months post-infection), including a complete loss of brain infiltrating CD8^+^ T cells (Fig. 5D-F). Strikingly, depleting T cells during virus chronicity dramatically improved cognitive performance as measured by Novel Object Recognition (NOR) and Discrimination Ratio behavioural analysis (DR) (Fig. 5B & C), suggesting that virus-induced T cell responses potentiate cognitive decline in this AD mouse model.

## Discussion

Elucidating the mechanisms through which viral infections promote AD development is essential to understand, predict and potentially mitigate the risk of infectious exacerbations of AD. Herein, using well-established mouse models of AD and herpesvirus infection, we identify that virus-induced T cell responses accelerate AD progression. We found that viral infection triggered an influx of T cells into the brain during AD that consisted primarily of clonally diverse virus-specific CD8^+^ T cells. Virus-induced cognitive decline in AD transgenic mice was ameliorated by administration of antiviral drugs which reduced brain T cell infiltrations, and by direct depletion of T cells. Thus, we demonstrate that a virus-induced immune response accelerates AD progression and provide evidence that this process could be therapeutically targeted for clinical benefit.

The mechanisms through which MCMV-induced T cells causes neuropathology are not fully understood. Given the clonal diverse nature of brain-infiltrating CD8^+^ T cells in virus-infected AD transgenic mice, together with detection of CD8^+^ T cell populations reactive to multiple virus-derived antigenic peptides, it is unlikely that virus-specific T cell responses are directly responding to and killing neurons. Interestingly, elevated concentrations of IFNγ have been observed in HCMV seropositive AD patients [23] which is expressed by leukocytes including virus-specific T cells in response to viral infections. IFNγ can inhibit dendritic outgrowth which associates with decreased rate of synapse formation [40]. Importantly, neutralization of IFNγ ameliorates pathology in a T cell-driven model of tauopathy [21] and reduces plaque burden in APP/PS1 mice [41]. Thus, these data imply that virus-induced T cells promote disease progression via the expression of IFNγ.

CD8^+^ T cells were the predominant brain-infiltrating T cell population that associated with infection-induced AD exacerbation and were detectable in a site of infection-induced neuronal damage, the hippocampus. The presence of CD8^+^ T cells in brains of AD patients has been described [17, 19] and EBV-specific CD8^+^, but not CD4^+^, T cells are detectable in the CSF of AD patients [18]. However, our data do not preclude a role for MCMV-induced CD4^+^ T cells in AD progression. Indeed, HLA-DRB1 is a risk-locus for AD [42], and CD4^+^ Th1 and Th17 cells exhibit inflammatory functionality in experimental AD models [41, 43]. Although CD4^+^ T cell were present at lower frequencies than CD8^+^ T cells in the brains of MCMV-infected 3xTg-AD, future work is required to disentangle specific roles for different T cell subsets in infection-induced AD progression.

Notably, 3xTg-AD mice exhibit splenomegaly and expansion of lymphoid cells in peripheral blood [37]. However, these peripheral dysregulations in immune responses in this model appeared not to dramatically influence viral-driven AD pathology in this model. Firstly, TCR deep sequencing analysis revealed similar TCR diversities in virus-specific CD8^+^ T cells in the periphery of both C57BL/6 and 3xTg-AD mice, suggesting that the virus-specific CD8^+^ T cells in the periphery were qualitatively similar in both models. Importantly, peripheral T cell perturbations in 3xTg-AD mice are dominated by CD4^-^CD8^-^CD3^+^ cells [37, 44], whereas deletion of both CD4^+^ and CD8^+^ T cells was required to ameliorate MCMV-driven AD acceleration. Thus, the dominant T cell type that is dysregulated in this AD model was shown to overtly contribute to viral-driven acceleration of AD progression.

The observation that a reduction but not a complete alleviation of virus-specific T cell responses by valaciclovir treatment was sufficient to ameliorate viral-driven AD exacerbation suggests that the degree of virus-specific T cell expansion rather than binary presence/absence of HCMV infection may be important in determining the influence of HCMV infection in AD progression. Given the dependence of epidemiological studies use of serology tests and associated binary outcomes, this may explain discordant results from epidemiological studies regarding the association between HCMV and AD. In the future, it will be important to investigate the relationship between virus-specific T cell expansions and AD risk.

Virus-induced exacerbation of AD development was not associated with detection of viral DNA in whole brain tissue. A limitation of our study is that analysis of whole-organ DNA may lack the sensitivity required to detect low levels of viral infection in specific brain regions such as the hippocampus. Thus, we cannot definitively conclude that MCMV was absent from the brain during AD development. Indeed, treatment of 3xTg-AD mice with antiviral drugs during the chronic phase of infection alleviated cognitive decline and associated T cell brain infiltrations. Notably, valaciclovir treatment impacts peripheral T cell responses during MCMV chronicity [28] which would impact T cell recruitment into brain tissue irrespective of presence of virus within the brain.

If MCMV is absent from brains of 3xTG-AD mice, how are virus-specific T cells recruited and activated within brain tissue? AD development is associated with an inflammatory environment within the brain that includes expression of T cell-attractant chemokines [45]. Thus, virus-specific T cells activated by peripheral viral infection could be recruited into the brain independently of virus replication. Given that IL-18, which was increased in brains of MCMV-infected 3xTg-AD mice and is increased in brains of AD patients [46], can promote T cell proliferation [47] and activate T cells in the absence of antigen [39], virus-specific T cells could be recruited, proliferate within brain tissue and be activated to produce cytokines and thus contribute to neuropathology in brains of AD patients irrespective of the neurotropism of the virus to which they have been primed by. Given that 45 different viruses and numerous other pathogens (including viruses that are not considered to be neurotropic) have been linked to increased risk of AD development [6] these data imply that infection-induced T cell responses may reflect a universal mechanism through which viral infections could contribute to the exacerbation of AD development (reviewed in: [16, 48]). Understanding the features of T cells that mediate viral exacerbation of AD and identifying mechanisms that impinge on these pathological responses will be critical for understanding and targeting viral-driven processes that worsen AD progression.

## Materials and Methods

### Ethics Statement

All mouse experiments were performed at Cardiff University and performed under the authority of UK Home Office project License (P8159A562).

### Mice and treatments

C57BL/6j WT mice (JAX stock #000664) were purchased from either Envigo or Charles River. 3xTg-AD mice contain three mutations associated with familial Alzheimer’s disease (APP Swedish, MAPT P301L, and PSEN1 M146V). They are bred in-house but were originally purchased from JAX laboratories (JAX stock #004807, B6;129-Tg(APPSwe,tauP301L)1Lfa *Psen1^tm1Mpm^*/Mmjax) [49]. Mice were infected with salivary gland-derived MCMV (Smith Strain) or mock infected with PBS i.p as described in figure legends. In some experiments, mice were treated with 1mg/ml Valaciclovir Hydrochloride (Cardiff and Vale NHS Pharmacy) dissolved in animal drinking water (replaced weekly) at day 60 post infection until experiment termination. In other experiments mice were treated with either 100μg of each anti-mouse anti-CD8 (clone YTS156.7.7 and YTS169.4) and anti-CD4 (clone GK1.5 and YTS191) depleting antibodies or isotype control (InVivoMAb rat IgG2b isotype control, anti-keyhole limpet hemocyanin (BioxCell) i.p at day 60 post infection, repeated twice, 7 days apart (400μg total per mouse).

### TCR deep sequencing

Twelve 8–10-week-old female C57BL/6 and 12 × 8–10-week-old female 3xTg-AD mice were infected with 5 × 10^4^ pfu MCMV i.p. At 9 months of age blood was extracted via cardiac puncture and animals were subsequently perfused with PBS and brains harvested. Leukocytes were extracted from both blood and brain (described below (cellular phenotyping)). Cells were pooled from 6 mice giving rise to 2 groups per phenotype (keeping tissue separate). Cells were stained with Zombie Aqua fixable viability dye (Biolegend), then stained with anti-CD16/CD32 Fc-block (Biolegend). Cells were next stained with 25 μg/ml MCMV-specific (H-2K^b^ M38 residues 316–323 (SSPPMFRV)) tetramer conjugated to PE for 15 minutes at 37 °C (monomer kindly provided by the National Institutes of Health Tetramer Core Facility). Cells were next stained with anti-CD8 APC (clone 53.6.7, Biolegend) and anti-CD3 BV605 (clone 145-2C11, Biolegend) (all anti-mouse), for 20 minutes at 4 °C. Data was acquired using a modified BD FACS ARIA (BD Biosciences) and CD8^+^/M38^+^ populations were sorted into RNA-protect buffer (Qiagen). Samples were shipped to iRepertoire for TCR deep-sequencing. RNA was extracted and amplified using Amp2seq (armPCR) using TCR-beta chain long-read libraries. Samples were sequenced using a Miseq Nano (Illumina) at a depth of approximately 125,000 reads per library.

### Animal behaviour

Animal behaviour and cognitive assessment was measured using novel object recognition (NOR) as described previously [50]. For each behavioural test a minimum of 4-10 mice were included per condition. Briefly, behavioural testing was performed using a custom-made square plastic test arena (39 cm (Height) × 39 cm (Width) × 39 cm (Length)) with three non-transparent walls and one transparent wall to observe animals. This was placed in a class II laminar flow hood affixed with a GoPro HERO session camera (GoPro, USA). Each animal is tested in isolation whereby each animal is placed in the box containing two identical objects (familiar objects, FO) for 10 mins. Each mouse was returned to its home cage for 20 mins post familiarisation. Finally, one FO was replaced with a novel object (NO) and each mouse was placed in the test arena for a further 10 min. The FOs and NOs were similar in size but differed in colour and shape. All recorded data was analysed using Ethovision XT 13 software (Noldus, Wageningen, The Netherlands). Discrimination ratios (DR) were determined by dividing the time spent exploring the NO (physical touching of object) by the time spent exploring both the FO and the NO. Impairment of cognition was shown by a DR below 0.5. The analyst was blinded to the group allocation during the test. Spontaneous alternation rate behaviour was performed via a T-maze test using a custom-made plastic T-shaped arena (Fig. 1D) with enclosed opaque sides as described previously [51]. Animals were not habituated to the test arena prior to testing. Initially the maze was set ensuring all partition doors are raised. The animal was placed into position A and is allowed to choose a goal arm (highlighted in red). Once a side is chosen the partition door was gently closed confining the animal for 30 seconds. The animal is then removed from the confined area (ensuring little stress) and the partition raised again. The animal was returned to position A facing away from the goal arms. The animal was then allowed to choose between the two open goal arms. Each test tool a minimum of 1 minute with 2 minutes per test on average. Each animal was assessed 10 times replacing the animal back into position A each time as described above. Alternation rate (AR) of 50% indicates the animal chose each open goal arm 50% of the time. The percentages indicate the number of times per 10 test per animal, that the animal chose the same goal arm.

### Viral load quantification

Organ specific infectious virus was quantified via plaque assay as described previously [52]. Viral DNA copy number in saliva was determined using qPCR for relative expression of IE1 using MCMV IE1 forward (50-AGCCACCAACATTGACCACGCAC-30) and MCMV IE1 reverse (50-GCCCCAACCAGGACACACAACTC30) primers, as described previously [53, 54].

### Cellular Phenotyping

For direct ex-vivo staining of blood, spleen and brain derived leukocytes from both C57BL/6 and 3xTg-AD AD mice, cells were stained as described previously [53, 55]. At experiment endpoint, blood was extracted via cardiac puncture and animals were subsequently perfused with PBS and brains harvested. Brains were cut into small pieces with dissection scissors and incubated with RPMI 1640 medium (Invitrogen) supplemented with 5 mM CaCl2, 5% FCS (Invitrogen), 1 mg/ml collagenase D (Roche Diagnostics), and 10 mg/ml DNAse I (Sigma) at 37°C for 45 minutes before passing through a 40 μM cell strainer. Cells were washed and separated using an additional Percoll (GE Healthcare) purification step. Cells were stained with Zombie Aqua fixable viability dye (Biolegend), then stained with anti-CD16/CD32 Fc-block (Biolegend) and either stained for tetramer expression and / or cell surface marker expression. Tetramer staining – MCMV-specific MHCI monomers were kindly provided by the National Institutes of Health Tetramer Core Facility. The biotinylated monomers were tetramerised described previously [53]. Cells were next stained with multiple combinations of cell surface expression markers (all anti-mouse) described below. Cell surface Marker expression - Cells were stained with various combinations of the following antibodies (all anti-mouse); anti-CD4 Brilliant-Violet 605 (BV605), PE/Cy7 or PerCp/Cy5.5 (clone RM4-5, Biolegend), anti-CD8 Brilliant-Violet 711 (BV711), Brilliant-Violet 605 (BV605), PerCp/Cy5.5, APC/Cy7 or PE/Cy7 (clone 53.6.7, Biolegend), anti-CD11a PerCp-Cy5.5 (clone M17/4, Biolegend), anti-CD44 FITC or APC/Cy7 (clone IM7, Biolegend), anti-CD45 Brilliant-Violet 605 (BV605) (clone 30-F11, Biolegend), anti-CD62L FITC, PE/Cy7 or BV711 (clone MEL-14, Biolegend), anti-CD69 PE/Cy7 or BV711 (clone H1.2F3, Biolegend), anti-CD103 PE/Dazzle (clone 2E7, Biolegend), for 20 minutes at for 20 minutes at 4 °C. Data were acquired using an Attune NxT flow cytometer (ThermoFisher Scientific). All flow cytometry data was analysed using FlowJo (Treestar).

### DiOlistic spine labelling

Freshly dissected and sectioned hippocampal slices (200 μm thickness, McIlwain tissue chopper) were analysed using DiOlistic labelling approach for the visualisation of the dendritic and synaptic architecture as previously described [56]. Briefly, tungsten particles (1.67 µm diameter, Bio-Rad) were coated with 1,1′-Dioctadecyl-3,3,3′,3′-Tetramethylindocarbocyanine Perchlorate (DiI) fluorometric dye (Life Tech) that had been pre-dissolved in dichloromethane. These particles were loaded in ethylene tetrafluoroethylene (ETFE) tubing and delivered onto the tissue slices at 100-120 psi using a Helios Gene Gun (Bio-Rad, United States) through a 3.0 μm pore size cell culture insert. Dye diffusion was facilitated for 20 mins in Neurobasal-A medium (Life Technologies) at 37°C with 5% CO_2_ and fixed with 4% paraformaldehyde (PFA) for 30 min at room temperature, nuclear stained with Hoechst 33342 and mounted in FluorSave. Dendritic spines on apical dendrites within the CA1 hippocampal neurons were imaged using the Leica SP8 laser-scanning confocal microscope with lightning deconvolution (Leica Microsystems, Germany). As described previously [56] dendritic segments longer than 30 μm were 3D reconstructed using the Filament Tracer module in Imaris software (v9.3.1, Bitplane) and spines subtyped with the Spine Classifier MATLAB extension. Predetermined settings were used to morphologically classify and subtype the spines based on the following rules; spines <0.8 μm: ‘stubby’ spines, between 0.8 and 3 μm but with a spine head diameter greater than neck width: ‘mushroom spines’; between 0.8 and 3 μm: ‘thin’ spines. Each reconstructed dendrite segment was manually checked to ensure correct spine detections. Each measurement was taken from at least four separate values from each mouse.

### Immunohistochemistry

Animals were perfused with PBS and brains were excised and fixed using 10% neutral-buffered formalin saline. Whole brains were paraffin embedded and 5 µm sections were cut from PBS or MCMV-infected C57BL/6 and 3xTg-AD mice. For fluorescent immunohistochemistry of immune cell infiltrate, sections were stained using an in-house method using a Leica Bond RX automated staining system with antigen retrieval performed using ER2 buffer (Leica Biosystems) prior to endogenous peroxidase activity quenching with 1% H_2_O_2_ (Sigma). Non-specific binding was blocked with 10% goat serum. Brain sections were incubated with anti-CD3 (mab ABclonal) and anti-CD8 (clone D4W2Z, Cell Signalling Technology) for 50 mins at room temperature (all anti-mouse). Bound primary antibody was detected with either anti-Rabbit VisUCyte (Bio-Techne) or anti-Rat ImmPRESS HRP polymer (Vector Labs) and next incubated with either 408nm, 488nm, and 594nm Tyramide conjugates (Biotium – conjugated in-house) for fluorescence signal generation before counterstaining with Hoechst 33342 (ThermoFisher). Slides were mounted using Vectashield Vibrance (Vector Labs) and imaged using Zeiss Axio Scan.Z1 slide scanner (Plan-Apochromat 20x/0.8NA) at 20 × magnification. The number of CD8^+^/CD3^+^ cells was quantified using positive cell detection in QuPath (version 0.5.1) (University of Edinburgh) [57].

### ELISA

Perfused brains (described above) were mechanically homogenized and supernatant was analysed via mouse IL-18 ELISA (Biolegend), human total Tau ELISA (Biolegend), human Tau (pT231) phospho ELISA (ThermoFisher), and human Aγ42 ELISA (Abcam) according to manufacturer protocols. Brain homogenate was further homogenized using Tris/EDTA buffer pH9 (Qiagen), 1% TritonX-100 (Sigma), 0.1% NaN3 and 1 × protease cocktail inhibitor set (Sigma) and analysed using human Aγ40 ELISA (ThernoFisher) according to manufacturer protocols.

### Statistics

Statistical analysis was performed using the Mann–Whitney U test or Welch’s t test for paired analysis or one-way and two-way ANOVA analysis with Tukey’s correction for multiple comparison tests. p values are reported as follows: n.s., >0.05; *, ≤0.05; **, ≤0.01; ***, ≤0.001; and ****, ≤0.0001. To assess biological replication: all in vivo experiments were performed multiple times as stated in figure legends. All outliers were included in datasets, as shown.

## Material availability

This study did not generate new unique reagents.

## Data and code availability

All original TCR-deep sequencing data will be publicly available as of the date of publication. Any additional information required to reanalyse the data reported in this paper is available from the lead contact upon request.

## Supplemental Figures

**Figure S1. 3xTg-AD mice do not have defective antiviral control.** (A) Weight loss and (B) viral genomes measured in saliva by qPCR at day 35 in C57BL/6 or 3xTg-AD mice infected with 5 × 10^4^ pfu MCMV (i.p.). Data are shown as mean ± SEM (A) or median (B) n = 10 C57BL/6 and 10 3xTg-AD mice per group representing five independent experiments. (C) MCMV genome load was assayed by qPCR detected of MCMV gB in brain homogenates of 9-10-month-old C57BL/6 or 3xTg-AD mice infected with 5 × 10^4^ pfu MCMV (i.p.) for 6-7 months or spleen homogenates of C57BL/6 infected with 5 × 10^4^ pfu MCMV (i.p.) 2 days post-infection with results from three independent replicates quantified relative to ACTIN. Data shown are representative of 2-5 independent experiments with 4 C57BL/6 and 10 3xTg-AD mice per group.

**Figure S2. Circulating CD8 T cells in MCMV-infected C57BL/6 and 3xTg-AD mice.** (A) Percentage expression of CD11a^+^/CD69 CD4^+^ tissue resident memory T cells (TRM) (left) and CD103^+^/CD69^+^ CD8^+^ TRM (right) quantified in brains via flow cytometry in MCMV-infected or non-infected C57BL/6 or 3xTg-AD mice. Some mice groups were treated with or without oral addition of 1mg/ml Valaciclovir Hydrochloride from day 60 post infection until experiment termination. (B-D) Percentage expression of (B) peripheral blood CD8^+^ cells and MCMV tetramer^+^ (C) IE3 and (D) M38 CD8^+^ T cells in MCMV-infected C57BL/6 or 3xTg-AD mice were quantified over a 180-day time-course via flow cytometry. (A-D) Data are shown as mean ± SEM; n = 4 C57BL/6 and 9-10 3xTg-AD mice per group representing two independent experiments).

**Figure S3. TCR diversity increases in the brains of MCMV-infected 3xTG-AD mice.** C57BL/6 or 3xTg-AD mice were infected with 5 × 10^4^ pfu MCMV (i.p.). Leukocytes were harvested from the brains and circulating blood at 6 months post-infection (9 months of age). (A) Concatenated frequency of murine TRBV (mTRBV) gene transcripts (key) and (B) normalized Shannon–Wiener index), reflecting clonality in relation to overall repertoire size of MCMV tetramer^+^ (M38) CD8^+^ T cells from the brain and blood of C57BL/6 or 3xTg-AD mice. All data shown are concatenated from M38-tetramer^+^ CD8^+^ T cells sorted from pooled C57BL/6 or 3xTg-AD mice (n = 5 per group, per tissue and is representative of two independent experiments).

**Figure S4. Increased viral-mediated pro-inflammatory cytokine release in AD susceptible mice.** (A) IL-18 concentration in brain homogenates of 9–10-month-old C57BL/6 or 3xTg-AD mice infected with 5 × 10^4^ pfu MCMV or PBS (i.p.) for 6-7 months measured by ELISA. Data shown are representative of 3-4 C57BL/6 and 10 3xTg-AD mice per group. Statistical significance was assessed using one-way ANOVA analysis with Tukey’s multiple comparisons test. p values are reported as follows: n.s., >0.05; *, ≤0.05; **, ≤0.01; ***, ≤0.001; and ****, ≤0.0001.

## Acknowledgments

The authors would like to thank Professor Andrew J Godkin (Cardiff University) for helpful discussion and constructive reading of the manuscript.

## References

1. Breijyeh, Z. and R. Karaman, Comprehensive Review on Alzheimer’s Disease: Causes and Treatment. Molecules, 2020. 25(24).

2. Sims, R., M. Hill, and J. Williams, The multiplex model of the genetics of Alzheimer’s disease. Nat Neurosci, 2020. 23(3): p. 311–322.

3. Fischer, O.L., Miliare Nekrosen mit drusigen Wucherungen der Neurofibrillen, eine regelmässige Veränderung der Hirnrinde bei seniler Demenz. Monatsschr Psychiatr Neurol, 1907. 22(4): p. 361–372.

4. Alzheimer, A., über eigenartige Krankheitsfälle des späteren Alters. Neur. u. Psych., 1911. 4: p. 356–385.

5. Vojtechova, I., et al., Infectious origin of Alzheimer’s disease: Amyloid beta as a component of brain antimicrobial immunity. PLoS Pathog, 2022. 18(11): p. e1010929.

6. Levine, K.S., et al., Virus exposure and neurodegenerative disease risk across national biobanks. Neuron, 2023. 111(7): p. 1086–1093 e2.

7. Wang, L., et al., Association of COVID-19 with New-Onset Alzheimer’s Disease. J Alzheimers Dis, 2022. 89(2): p. 411–414.

8. Wozniak, M.A., A.P. Mee, and R.F. Itzhaki, Herpes simplex virus type 1 DNA is located within Alzheimer’s disease amyloid plaques. J Pathol, 2009. 217(1): p. 131–8.

9. Jamieson, G.A., et al., Herpes simplex virus type 1 DNA is present in specific regions of brain from aged people with and without senile dementia of the Alzheimer type. J Pathol, 1992. 167(4): p. 365–8.

10. Readhead, B., et al., Multiscale Analysis of Independent Alzheimer’s Cohorts Finds Disruption of Molecular, Genetic, and Clinical Networks by Human Herpesvirus. Neuron, 2018. 99(1): p. 64–82 e7.

11. Taquet, M., et al., The recombinant shingles vaccine is associated with lower risk of dementia. Nat Med, 2024. 30(10): p. 2777–2781.

12. Barnes, L.L., et al., Cytomegalovirus infection and risk of Alzheimer disease in older black and white individuals. J Infect Dis, 2015. 211(2): p. 230–7.

13. Aiello, A.E., et al., The influence of latent viral infection on rate of cognitive decline over 4 years. J Am Geriatr Soc, 2006. 54(7): p. 1046–54.

14. Readhead, B.P., et al., Alzheimer’s disease-associated CD83(+) microglia are linked with increased immunoglobulin G4 and human cytomegalovirus in the gut, vagal nerve, and brain. Alzheimers Dement, 2025. 21(1): p. e14401.

15. Wennberg, A.M., et al., Association of common infections with cognitive performance in the Baltimore Epidemiologic Catchment Area study follow-up. Alzheimers Dement, 2023. 19(11): p. 4841–4851.

16. Clement, M., The association of microbial infection and adaptive immune cell activation in Alzheimer’s disease. Discov Immunol, 2023. 2(1): p. kyad015.

17. Merlini, M., et al., Extravascular CD3+ T Cells in Brains of Alzheimer Disease Patients Correlate with Tau but Not with Amyloid Pathology: An Immunohistochemical Study. Neurodegener Dis, 2018. 18(1): p. 49–56.

18. Gate, D., et al., Clonally expanded CD8 T cells patrol the cerebrospinal fluid in Alzheimer’s disease. Nature, 2020. 577(7790): p. 399–404.

19. Unger, M.S., et al., CD8(+) T-cells infiltrate Alzheimer’s disease brains and regulate neuronal- and synapse-related gene expression in APP-PS1 transgenic mice. Brain Behav Immun, 2020. 89: p. 67–86.

20. Lee, S.H., et al., TREM2-independent oligodendrocyte, astrocyte, and T cell responses to tau and amyloid pathology in mouse models of Alzheimer disease. Cell Rep, 2021. 37(13): p. 110158.

21. Chen, X., et al., Microglia-mediated T cell infiltration drives neurodegeneration in tauopathy. Nature, 2023. 615(7953): p. 668–677.

22. Dai, L. and Y. Shen, Insights into T-cell dysfunction in Alzheimer’s disease. Aging Cell, 2021. 20(12): p. e13511.

23. Lurain, N.S., et al., Virological and immunological characteristics of human cytomegalovirus infection associated with Alzheimer disease. J Infect Dis, 2013. 208(4): p. 564–72.

24. Westman, G., et al., Increased inflammatory response in cytomegalovirus seropositive patients with Alzheimer’s disease. PLoS One, 2014. 9(5): p. e96779.

25. Harrison, M.A.A., et al., Intermittent cytomegalovirus infection alters neurobiological metabolism and induces cognitive deficits in mice. Brain Behav Immun, 2024. 117: p. 36–50.

26. Reuter, J.D., et al., Systemic immune deficiency necessary for cytomegalovirus invasion of the mature brain. J Virol, 2004. 78(3): p. 1473–87.

27. Krstanovic, F., et al., Susceptibility of Mouse Brain to MCMV Infection and Neuroinflammation During Ontogeny. Pathogens, 2024. 13(12).

28. Beswick, M., et al., Antiviral therapy can reverse the development of immune senescence in elderly mice with latent cytomegalovirus infection. J Virol, 2013. 87(2): p. 779–89.

29. Pchitskaya, E. and I. Bezprozvanny, Dendritic Spines Shape Analysis-Classification or Clusterization? Perspective. Front Synaptic Neurosci, 2020. 12: p. 31.

30. Helm, M.S., et al., A large-scale nanoscopy and biochemistry analysis of postsynaptic dendritic spines. Nat Neurosci, 2021. 24(8): p. 1151–1162.

31. Javonillo, D.I., et al., Systematic Phenotyping and Characterization of the 3xTg-AD Mouse Model of Alzheimer’s Disease. Front Neurosci, 2021. 15: p. 785276.

32. Mody, P.H., et al., Cytomegalovirus infection induces Alzheimer’s disease-associated alterations in tau. J Neurovirol, 2023. 29(4): p. 400–415.

33. Munks, M.W., et al., Genome-wide analysis reveals a highly diverse CD8 T cell response to murine cytomegalovirus. J Immunol, 2006. 176(6): p. 3760–6.

34. Arens, R., et al., Cutting edge: murine cytomegalovirus induces a polyfunctional CD4 T cell response. J Immunol, 2008. 180(10): p. 6472–6.

35. Walton, S.M., et al., The dynamics of mouse cytomegalovirus-specific CD4 T cell responses during acute and latent infection. J Immunol, 2008. 181(2): p. 1128–34.

36. Turner, S.J., et al., Structural determinants of T-cell receptor bias in immunity. Nat Rev Immunol, 2006. 6(12): p. 883–94.

37. Nava Catorce, M., G. Acero, and G. Gevorkian, Age- and sex-dependent alterations in the peripheral immune system in the 3xTg-AD mouse model of Alzheimer’s disease: Increased proportion of CD3+CD4-CD8-double-negative T cells in the blood. J Neuroimmunol, 2021. 360: p. 577720.

38. Venturi, V., et al., Sharing of T cell receptors in antigen-specific responses is driven by convergent recombination. Proc Natl Acad Sci U S A, 2006. 103(49): p. 18691–6.

39. Berg, R.E., et al., Memory CD8+ T cells provide innate immune protection against Listeria monocytogenes in the absence of cognate antigen. J Exp Med, 2003. 198(10): p. 1583–93.

40. Kim, I.J., et al., Interferon gamma induces retrograde dendritic retraction and inhibits synapse formation. J Neurosci, 2002. 22(11): p. 4530–9.

41. Browne, T.C., et al., IFN-gamma Production by amyloid beta-specific Th1 cells promotes microglial activation and increases plaque burden in a mouse model of Alzheimer’s disease. J Immunol, 2013. 190(5): p. 2241–51.

42. Kunkle, B.W., et al., Author Correction: Genetic meta-analysis of diagnosed Alzheimer’s disease identifies new risk loci and implicates Abeta, tau, immunity and lipid processing. Nat Genet, 2019. 51(9): p. 1423–1424.

43. Machhi, J., et al., CD4+ effector T cells accelerate Alzheimer’s disease in mice. J Neuroinflammation, 2021. 18(1): p. 272.

44. Marchese, M., et al., Autoimmune manifestations in the 3xTg-AD model of Alzheimer’s disease. J Alzheimers Dis, 2014. 39(1): p. 191–210.

45. Wojcieszak, J., K. Kuczynska, and J.B. Zawilska, Role of Chemokines in the Development and Progression of Alzheimer’s Disease. J Mol Neurosci, 2022. 72(9): p. 1929–1951.

46. Ojala, J., et al., Expression of interleukin-18 is increased in the brains of Alzheimer’s disease patients. Neurobiol Aging, 2009. 30(2): p. 198–209.

47. Iwai, Y., et al., An IFN-gamma-IL-18 signaling loop accelerates memory CD8+ T cell proliferation. PLoS One, 2008. 3(6): p. e2404.

48. Hosseininasab, S.S.M., et al., Alzheimer’s disease and infectious agents: a comprehensive review of pathogenic mechanisms and microRNA roles. Front Neurosci, 2024. 18: p. 1513095.

49. Oddo, S., et al., Triple-transgenic model of Alzheimer’s disease with plaques and tangles: intracellular Abeta and synaptic dysfunction. Neuron, 2003. 39(3): p. 409–21.

50. Webberley, T.S., et al., Assessment of Lab4P Probiotic Effects on Cognition in 3xTg-AD Alzheimer’s Disease Model Mice and the SH-SY5Y Neuronal Cell Line. Int J Mol Sci, 2023. 24(5).

51. Deacon, R.M. and J.N. Rawlins, T-maze alternation in the rodent. Nat Protoc, 2006. 1(1): p. 7–12.

52. Stack, G., et al., CD200 receptor restriction of myeloid cell responses antagonizes antiviral immunity and facilitates cytomegalovirus persistence within mucosal tissue. PLoS Pathog, 2015. 11(2): p. e1004641.

53. Clement, M., et al., Cytomegalovirus-Specific IL-10-Producing CD4+ T Cells Are Governed by Type-I IFN-Induced IL-27 and Promote Virus Persistence. PLoS Pathog, 2016. 12(12): p. e1006050.

54. Kamimura, Y. and L.L. Lanier, Rapid and sequential quantitation of salivary gland-associated mouse cytomegalovirus in oral lavage. J Virol Methods, 2014. 205: p. 53–6.

55. Clement, M., et al., Inhibitory IL-10-producing CD4(+) T cells are T-bet-dependent and facilitate cytomegalovirus persistence via coexpression of arginase-1. Elife, 2023. 12.

56. Bevan, R.J., et al., Improved DiOlistic labelling technique for neurons in situ: Detailed visualisation of dendritic spines and concurrent histochemical-detection in fixed tissue. Methods, 2024. 229: p. 82–93.

57. Bankhead, P., et al., QuPath: Open source software for digital pathology image analysis. Sci Rep, 2017. 7(1): p. 16878.

