## Supplementary figures and images for "Cytomegalovirus-induced T cell responses accelerate Alzheimer’s disease progression in mice"

### Supplemental Figure 1

#### Supplmental Figure 1

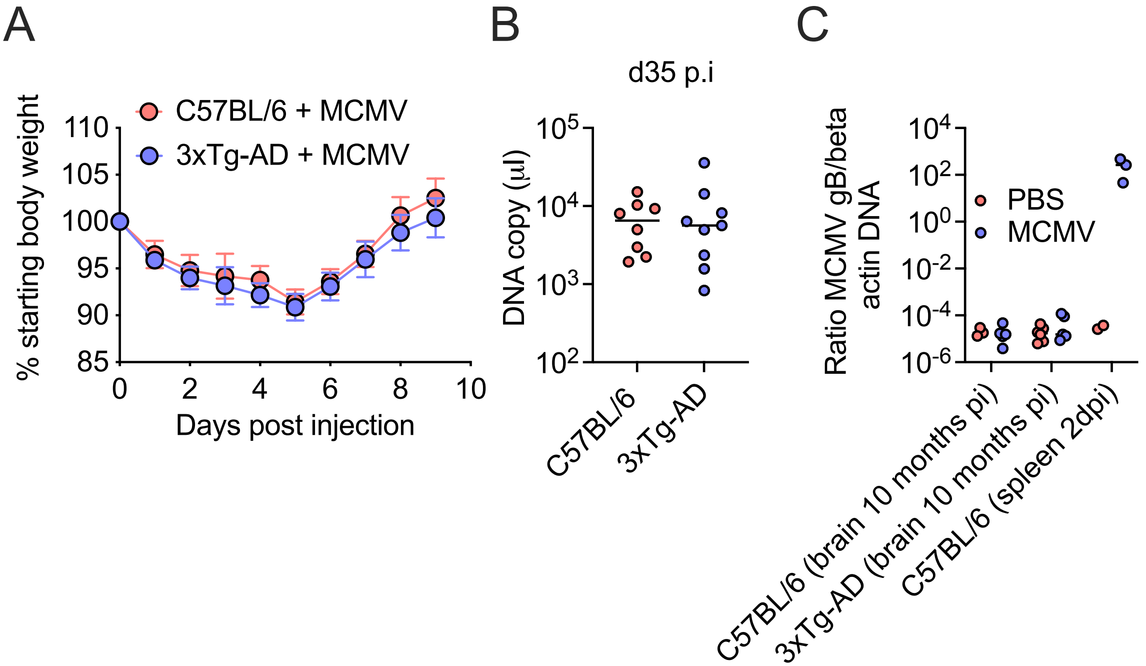

### Supplemental Figure 2

#### Supplmental Figure 2

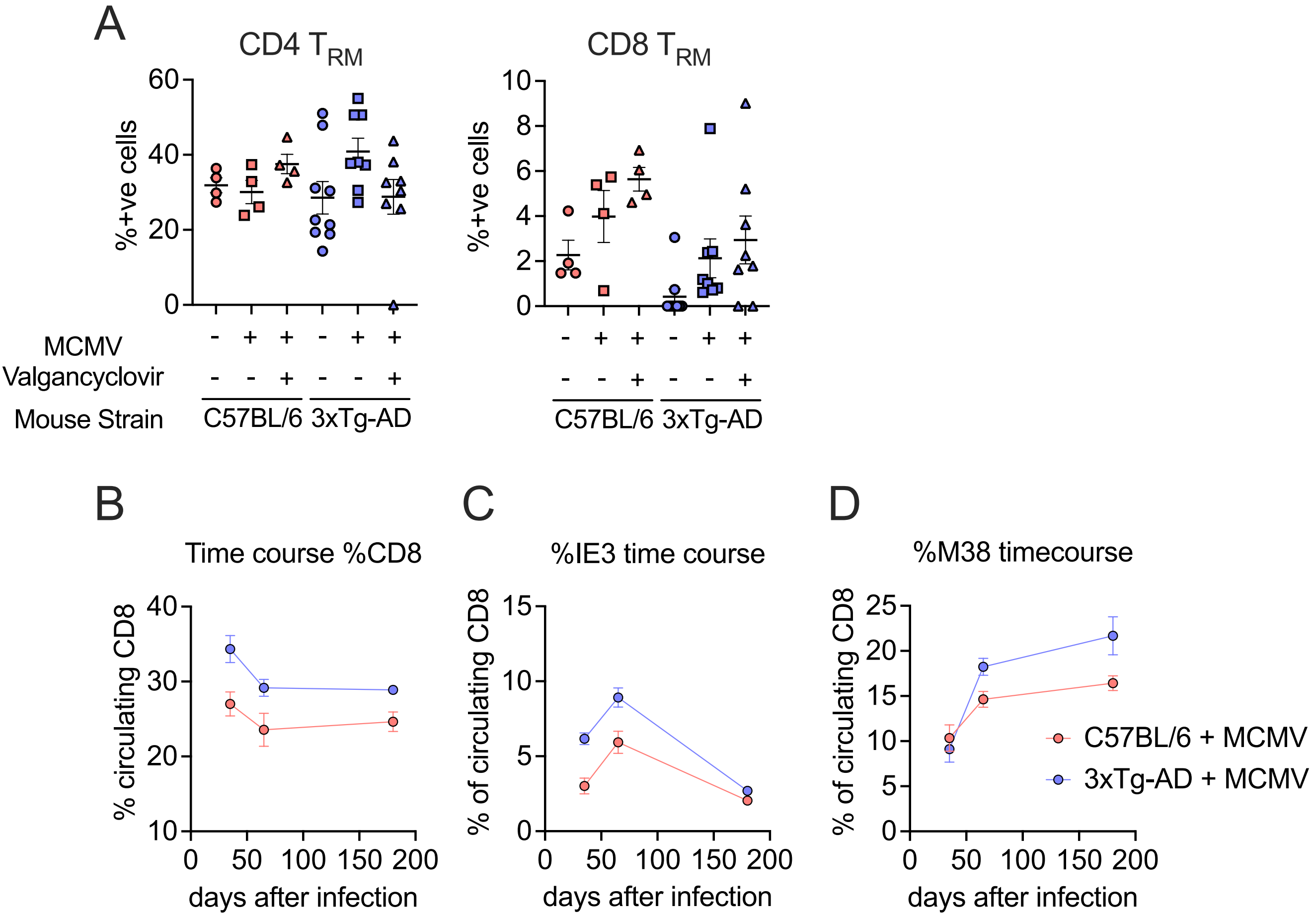

### Supplemental Figure 3

#### Supplemental Figure 3

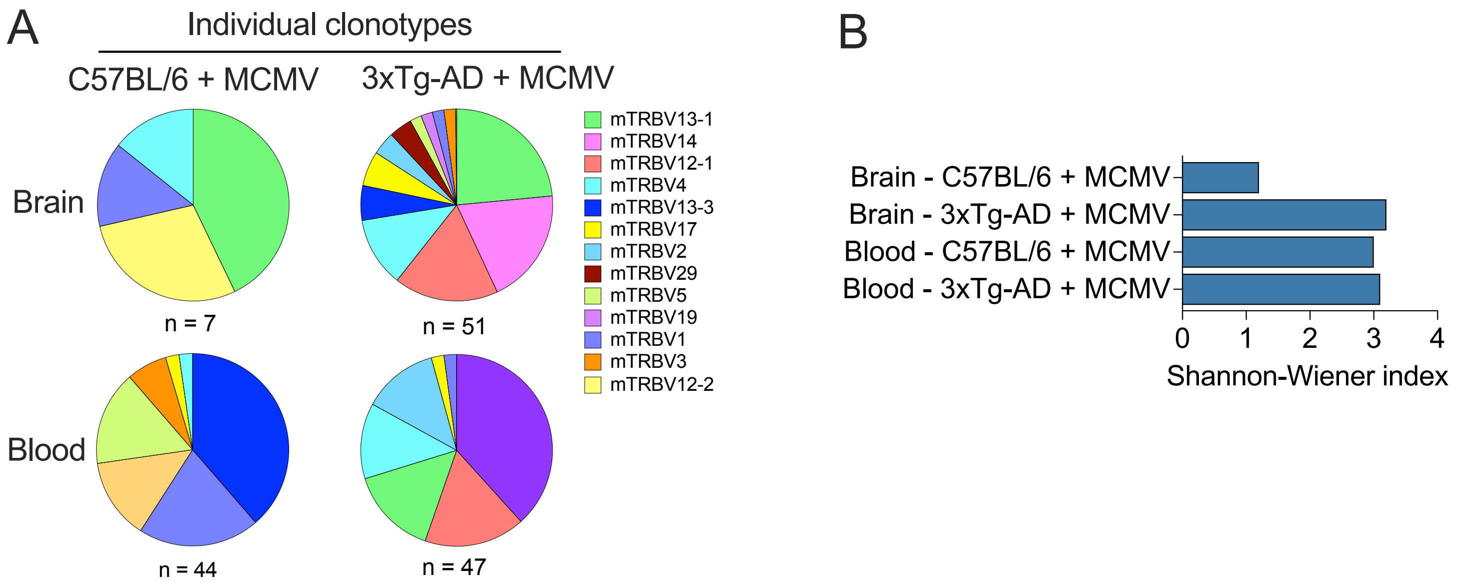

### Supplemental Figure 4

#### Supplemental Figure 4

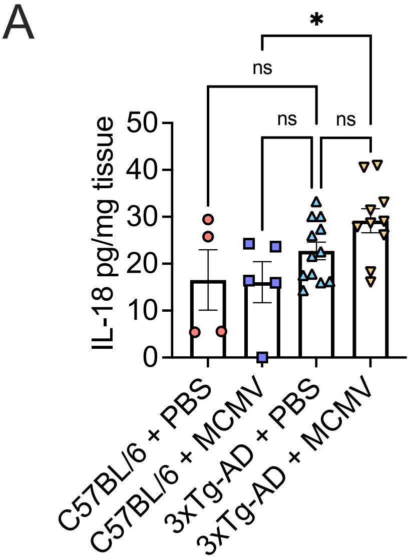
